# NeuroRoots, a bio-inspired, seamless Brain Machine Interface device for long-term recording

**DOI:** 10.1101/460949

**Authors:** Marc D. Ferro, Christopher M. Proctor, Alexander Gonzalez, Eric Zhao, Andrea Slezia, Jolien Pas, Gerwin Dijk, Mary J. Donahue, Adam Williamson, Georges G. Malliaras, Lisa Giocomo, Nicholas A. Melosh

## Abstract

Minimally invasive electrodes of cellular scale that approach a bio-integrative level of neural recording could enable the development of scalable brain machine interfaces that stably interface with the same neural populations over long period of time.

In this paper, we designed and created NeuroRoots, a bio-mimetic multi-channel implant sharing similar dimension (10µm wide, 1.5µm thick), mechanical flexibility and spatial distribution as axon bundles in the brain. A simple approach of delivery is reported based on the assembly and controllable immobilization of the electrode onto a 35µm microwire shuttle by using capillarity and surface-tension in aqueous solution. Once implanted into targeted regions of the brain, the microwire was retracted leaving NeuroRoots in the biological tissue with minimal surgical footprint and perturbation of existing neural architectures within the tissue. NeuroRoots was implanted using a platform compatible with commercially available electrophysiology rigs and with measurements of interests in behavioral experiments in adult rats freely moving into maze. We demonstrated that NeuroRoots electrodes reliably detected action potentials for at least 7 weeks and the signal amplitude and shape remained relatively constant during long-term implantation.

This research represents a step forward in the direction of developing the next generation of seamless brain-machine interface to study and modulate the activities of specific sub-populations of neurons, and to develop therapies for a plethora of neurological diseases.

## Introduction

Brain machine interfaces (BMIs) are playing an increasingly important role in neurological research (*1*–*4*), clinical treatments (*5*, *6*) and neural-prosthetics (*7*–*9*). With the advent of powerful signal processing and data analytics software tools, the impetus has shifted from two-dimensional bulky probes to developing electrical probes with higher channel counts, lower tissue damage, and long-term recording stability at the single cell level. Moreover, as these advances enable the simultaneous exploration of different part of the brain and potential clinical applications, straight-forward implantation strategies and minimal footprint have become an essential part of the implant development. Previous generations of bulky, stiff electrodes are being replaced by a number of innovative new devices (*10*, *11*) for lowering tissue damage (*12*), ultra-flexible meshes with chronic stability (*13*, *14*), flexible arrays (*2*, *15*, *16*), high channel counts (*17*, *18*) and stretchable electrodes (*3*, *19*). However, achieving the optimal combination of low tissue damage, scalability and facile surgical implantation remains challenging.

Inherent in these planar devices is that the electrode recording sites are tiled on a two-dimensional surface, restricting neuronal sampling to the volume directly in contact with this plane. This restricts the number of sites possible at a given depth and location. Increasing the number of electrodes on this plane by shrinking device dimensions results in higher resolution sampling of the neurons next to the shank but may not increase access to a higher number of unique neurons. This can be overcome by making wider electrode arrays which are in contact with more neurons at a given depth, yet these are in turn more mechanically disruptive. Interestingly, recent studies have found that porous planar electrode, with an open ‘mesh’ design, have low amounts of long-term tissue damage (*14*). However, because the mesh is still a continuous plane, delivery necessitates large needles and disruption of the initial neural structure, with a concomitant extended recovery period while neurons re-populate the insertion area.

To overcome the limitations of planar designs, we sought inspiration from biological structures. Communication between different regions in the brain is coordinated through of bundles of myelinated axons, making up the white matter within the brain. Human cortical axons range from 0.5-9 um in diameter (*20*), and can be many centimeters in length, with elastic moduli of roughly 10 kPa (*21*).

Here, we present axonal bundle mimics with similar spatial distribution and design of natural axonal architectures. These “NeuroRoots” (Fig 1) consist of arrays of individual electrodes, ∼7 um wide, ∼1.5 um thick, yet centimeters long, organized in axon-like tendrils. Each electrode has a single, ∼10 um diameter recording pad at its tip, and is mechanically separate from the other electrodes. This is a significantly different design from arrays of electrodes on a single shank, where the device width must increase to accommodate more electrodes at the same location. Instead, each electrode is independent, allowing complete flexibility in the number of electrodes at a given depth, without increasing the electrode width, and thus damage. Moreover, the electrodes have similar mechanical flexibility as myelinated axons, ideally enhancing long-term stability of the electrodes while lowering immunogenicity.

**Figure 1.**
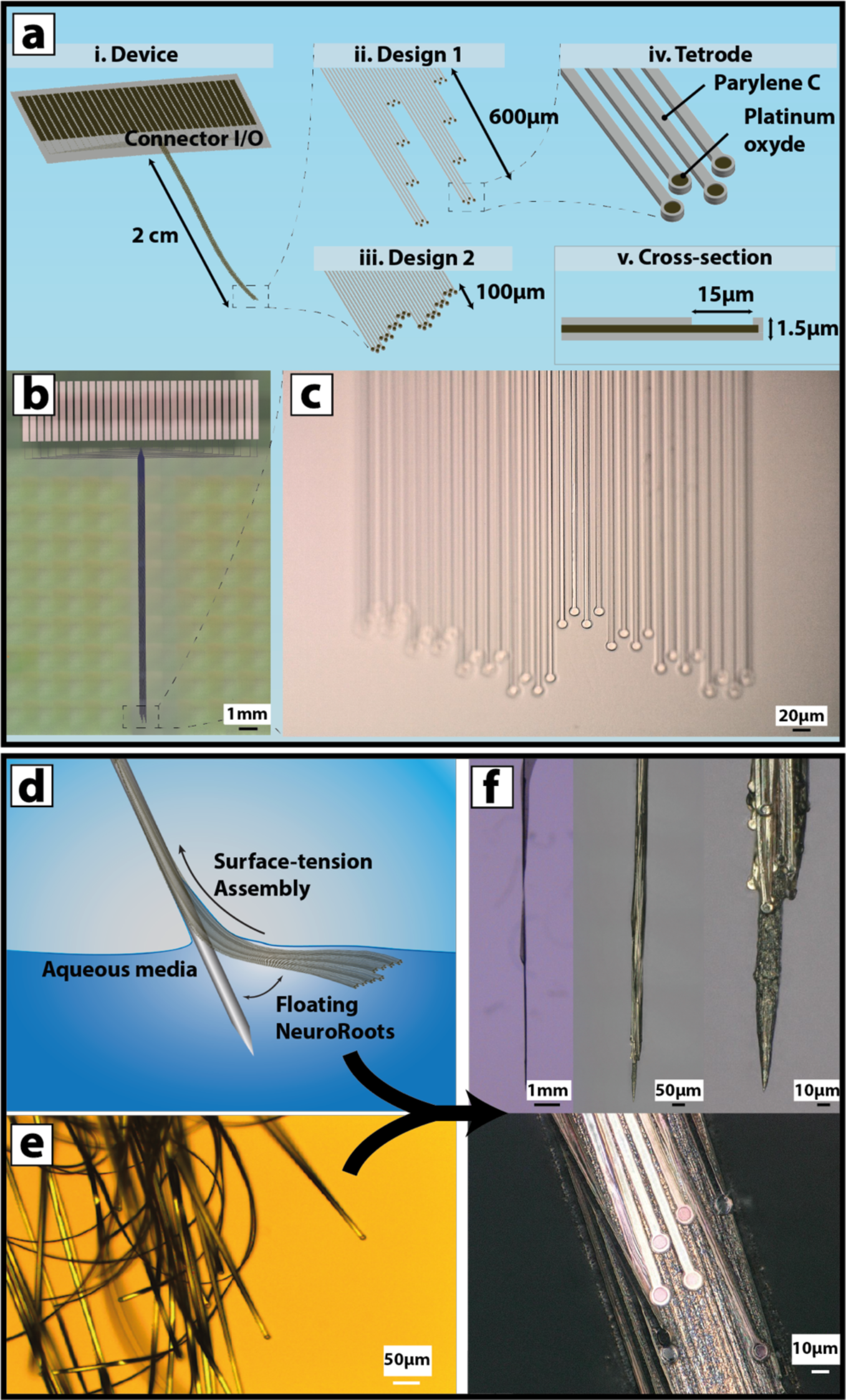
NeuroRoots overview and assembly. a) 3D model of the device and different configuration. i) NeuroRoots overview ii) Zoomed-in representation of the tip with the electrodes organized in 150µm depth spacing iii) representation of the 25µm depth spacing iv) Zoomed-in representation of one tetrode-like set of electrode v) Cross-sectional representation of one NeuroRoots electrode. The Parylene C substrate is represented in light grey and the platinum in dark brown. b) Microscope picture image of the NeuroRoots with Design 1 and Design 2, (c). d) Assembly method using capillary and surface-tension effects used to draw the electrodes onto the microwire. e) Microscope picture of the electrode leads after delamination from the fabrication substrate. f) Microscopy showing the electrodes assembled onto the microwires, demonstrating their spacing is set by the initial position in the array.

A critical challenge to these and other very soft electrodes is insertion into the brain due to their fragility and lack of mechanical stiffness. Previous research has shown that compliant electrodes can be inserted using mechanical shuttles (*22*), stiffening agents (*23*, *24*), or syringes (*14*). However, standard shuttles for a planar array of the NeuroRoots would be hundreds of microns wide and cause significant damage on their own. Instead, we developed an electrode self-assembly method using capillarity to organize large arrays of NeuroRoots onto a microwire as small as 20µm in diameter, which in turn provides mechanical support to allow implantation, yet with minimal damage (*25*, *26*). After insertion, the microwire could be removed leaving the electrodes in the tissue with the recording sites distributed according to their original length.

The microwire insertion platform is particularly convenient, as it enables direct use of traditional tetrode surgical apparatus for spatial targeting, implantation, and data acquisition. Avoiding complex, bulky connectorization and surgery significantly increased usability. After assembly onto the microwire, NeuroRoots were inserted into rat hippocampus using a standard Neuralynx Halo tetrode apparatus (*27*) and demonstrated stable recordings of action potentials in a freely-moving rat over seven weeks. Without further adjustments or calibration, the shape and amplitude of the signal recorded minimally changed over the time of the experiment, demonstrating long-term recording stability, with a biomimetic spatial distribution of recording sites. The combination of scalability, low damage, stable single unit recording, and ready integration with existing surgical apparatus make NeuroRoots a promising candidate for basic neuroscience experiments and clinical applications.

## Results

### Fabrication and Electrical Performance

The NeuroRoots design (Fig. 1.a) consists of independent polymer/metal/polymer ‘roots’, with thin leads connecting between the exposed recording pads at the tips, and larger pads at the proximal end that can be connected to standard electrophysiological acquisition systems. The specific device sizes, number of electrodes, and electrode materials were readily varied using standard photolithography and etching techniques. Parylene C, a flexible and biocompatible polymer, was chosen as substrate and insulator to encapsulate the Pt film used as conductive layer. In a typical preparation, 0.75 µm of Parylene-C was deposited onto a wafer and 5 µm wide, 100 nm thick Pt leads where then patterned via deposited and lift-off onto the Parylene-C. An additional 0.75 µm thick Parylene-C insulating layer was deposited on top (Fig 1b) and the outline of the device was photopatterned into 7 µm wide leads with 10 or 15 µm circular electrode pads at the end (Fig 1), and the excess material etched away in oxygen plasma. The total thickness of the device was measured at 1.5 µm (Fig. 1.a Cross-section). Parylene C is about 40 times stiffer than a human axon with a Young modulus of 400 kPa, though the geometrical thickness of NeuroRoots exhibit a bending stiffness equivalent to a human axon of 9 µm in diameter.

At the end of each lead was a circular electrode recording pad with a window in the upper Parylene layer to expose the bare Pt metal. The small electrodes configuration was organized into clusters of tetrodes (Fig. 1.a). The spatial distribution of these leads could be varied depending on the desired locations of the electrodes in the tissue, as the insertion process preserved the relative position of each electrode. Figure 1.a shows the different organizations of the 32 electrodes at the tip of the implants, either densely packed into layer of 100 µm in longitudinal depth (Fig. 1.c) or distributed over a 600 µm deep layer (Fig. 1.b). In an alternative design, the electrodes were coated with PEDOT:PSS and had a rectangular shape the width of the lead with a length up to 100 µm. The rough Pt or PEDOT-PSS pads at each tip provided low impedance electrodes. For example, the 15 µm diameter Pt electrodes exhibited an average impedance of 40 kΩ at 1 kHz, i.e. a specific impedance of 55 Ω.µm (Fig. S1). This is more than an order of magnitude lower than the impedance of smooth noble metal electrodes, and comparable to electrode of similar surface-area coated with a thin layer of PEDOT:PSS. As a direct comparison, a typical wire used in a tetrode exhibit the same electrode surface area but an average impedance of 300 kΩ. The dimensions of the electrodes together with their low impedance provide a superior platform for the low-noise recording of localized electrophysiological activity.

### Assembly

In order to insert electrodes with such extreme flexibility and miniaturization, a method of self-assembly was developed to controllably immobilize the roots of the implant onto a microwire that provides enough mechanical rigidity to penetrate the brain tissue using a tetrode surgical apparatus. (Fig. 1.g).

The first step of this process was to dip the NeuroRoots in a deionized water solution at the end of the microfabrication process which allowed the film to detach from the substrate and float at the air/liquid interface. Parylene-C is well-known for its hydrophobicity (*28*), and thus exhibits a high interfacial energy with water which allowed the film to unfold and float in its initial configuration.

The second step was to bring a tungsten microwire in contact with the implant and lift the floating electrodes onto the microwire substrate (Fig. 1d). By capillarity, a small amount of the aqueous solution would coat the microwire and offer a preferred energetic solution. Consequently the floating electrodes could be transferred onto a variety of materials and devices, including flexible plastics and inorganic shaped silicon (*29*). Here, the microwire was immersed into the water at an approximately 45° angle and the electrodes were subsequently transferred onto the microwire by withdrawing it from the water, allowing surface tension to draw the NeuroRoots onto the microwire surface (Fig. 1.e). Critically, capillary forces and surface tension directed all of the individual leads of the NeuroRoots to self-assemble along the length of the microwire despite the fact that initial configuration of the leads covered nearly ten times the width of the microwire. The floating electrodes were handled by the connector I/O section, which avoided damaging the roots themselves and offered a macroscopic handle to the implant.

In order to minimize the footprint and invasiveness, we used microwires as small as 35 µm diameter electrosharpened at the tip to a measured diameter of 20µm down to a few 100 nm (Fig. 1.g). Once assembled, the electrode bundle is less than the size of a single tetrode, yet with 8 times the recording capacity. The cross-sectional dimension is more than half the size compared to a single Utah array shank (80µm in diameter, (*30*)), a Michigan standard probe (125 to 50µm (*31*)), or even the recent achievement of ultra-thin silicon Neuropixel probe (70wide x20µm thick, with 100 recording sites per mm, (*17*)). (Fig. 1.g)

### Implantation

Different strategies have been previously explored in order to deliver minimal profile electronics into deep brain regions. These range from temporary attachments to a stiff micro-fabricated backing (*22*), molding of a rigid shanks using bioresorbable polymers (*23*), or more recently, temporary engaging onto a rigid carbon fiber (*12*) or injection using a standard needle (*32*). However, all those strategies present inherent limitations such as high initial insertion injury, limited scalability of electrode count, and complicated implantation procedures.

Here, our strategy was to add a small concentration of bio-soluble, inert polymer in the aqueous solution used during the assembly of the NeuroRoots so that we could immobilize the electrodes onto the microwire for a controllable period of time. We found that using a mixture of low molecular weight Polyethylene Glycol (PEG) in DI water could provide a release time frame between 5 to 20 minutes depending on the PEG concentration.

Once inserted into the brain region of interest, the microwire was retracted, leaving the roots innervated into the neural tissue (Fig. 2.c and d). Using this approach, we could accurately implant the electrodes into the brain with minimal surgical footprint, preventing large disruption of the Blood Brain Barrier (BBB) and minimizing local bleeding which is crucial in order to mitigate both initial damage and chronic tissue inflammation usually seen with mechanically rigid implant (*33*). In order to assess the maximal damage that could be done to the tissue using this approach and to image the NeuroRoots using the maximal resolution available with our *in-vivo* X-ray microtomography (µCT), we used the largest PEDOT:PSS implants. Histology performed on 2 rats and 2 mice did not reveal an important activation of microglia and reactive astrocyte around the implants (Fig. S2). We noticed when imaging chronic histology of smaller feature implants that a large portion of neural tissue often came off when the implant was removed, indicating that the NeuroRoots are able to tightly innervate the surrounding tissue (Fig. 2.d), corroborated by the stable chronic recordings discussed later.

**Figure 2:**
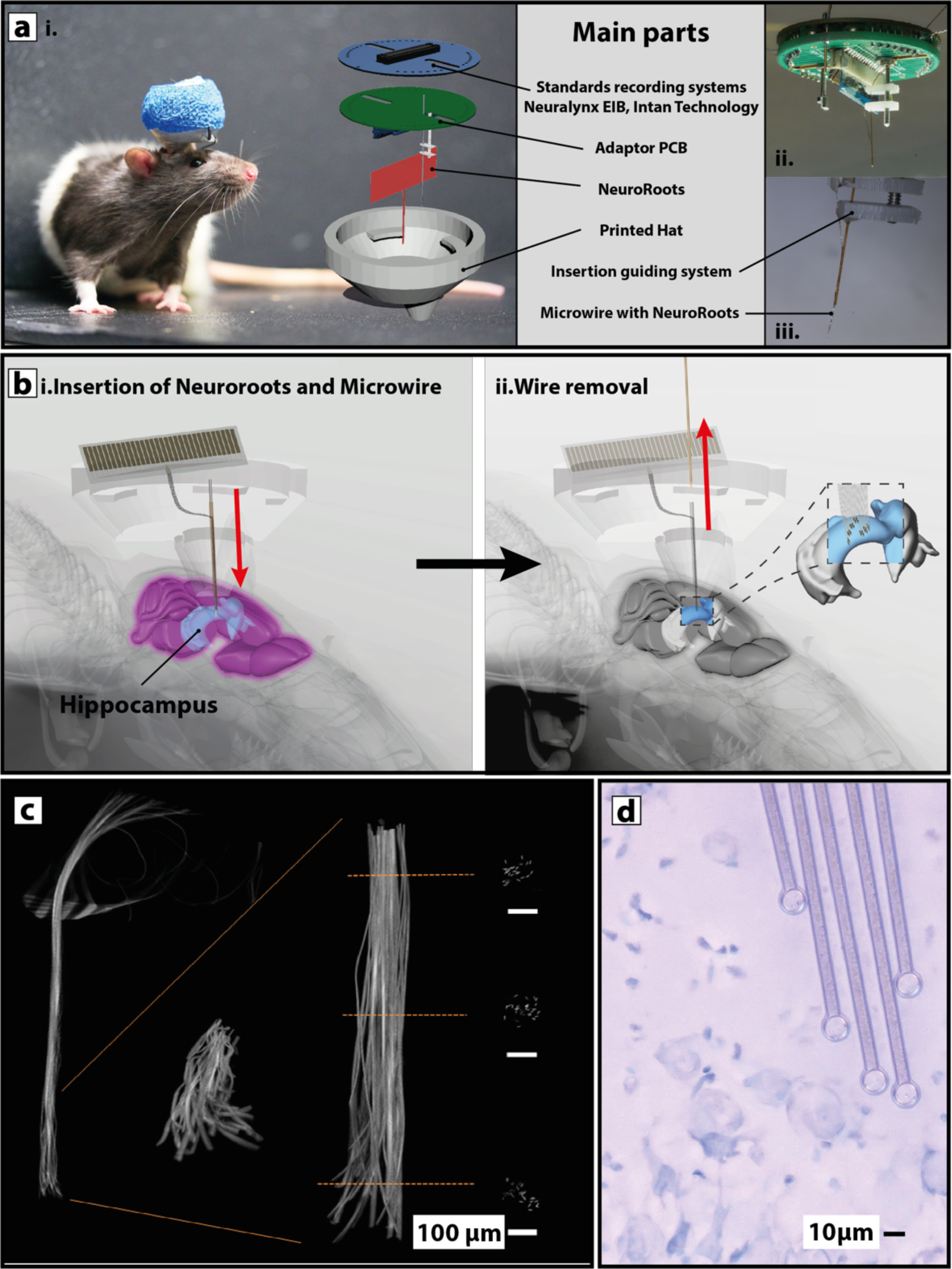
a) Apparatus for chronic recordings in freely moving rats. i) Picture of a rat 4 days post-surgery. The implant connector is protected by a cap on top of the scaffold presented on the right. The 3D exploded-view shows the main parts that compose the NeuroRoots platform. ii) Adaptor and the NeuroRoots assembled before insertion iii) Zoomed-in picture of the guiding system with the NeuroRoots assembled onto the microwire. b) Implantation strategy of the NeuroRoots into deep-brain regions. i) The assembled microwire and NeuroRoots are implanted into the desired brain region through a 3d printed scaffold hat. ii) once the NeuroRoots are released, the microwire is removed and leaving only the electrodes implanted into the brain tissue. c) X-ray microtomography scanning showing the electrode distribution after implantation using PEDOT:PSS based devices. d) Microscope image of a NeuroRoots placed onto a brain slice of cell the CA1 of the hippocampus for scale. The electrodes of 10 µm in diameter are similar in size with neuron soma. (Crecyl Violet staining)

An additional advantage of this implantation strategy is the reduced risk of mechanical failure after insertion compared with rigid implants for which can account for 50% of all failure modes (*34*). After the microwire removal, the implant was provided with additional mechanical slack by lowering the Z axis of the stereotaxic frame about 500µm before sealing the implant to the skull. This allowed for decoupling of the direct mechanical constrain between the implant and the brain which is under constant micromotion (*35*). Using this approach, we did not register any acute or chronic mechanical failure for the NeuroRoots.

### Apparatus

We demonstrated chronic recording of the NeuroRoots in fully mature rats freely moving in a complex maze equipped with infrared video tracking and automated reward systems (Fig. 2.a.i).

A Neuralynx ‘Halo 18’ was used as the surgical platform to make the system compatible with commercially available electrophysiology and behavioral rigs, thus minimally impacting the surgery procedures or protocols. A guide system was engineered to interface the NeuroRoots microelectrode and enable a precise alignment of the microwire, compatible with targeting using standard stereotaxic approach (Fig. 2 a.ii and a.iii)).

The NeuroRoots were then connected to an Intan Technology digital amplifier using a Zero Insertion Force (ZIF) connector. This then mated to a custom design Printed Circuit Board (PCB) to an Omnetics 36 channel connector or the EIB 72 of Neuralynx to the amplifier. (Fig. 2.a.ii). The entire platform was then securely assembled into a 3D-printed scaffold hat with only the tip of the implant protruding. The weight of the final device was measured to be 8 g, which is less than 2% of an adult rat weight and below the 10% bar recommended by *in-vivo* guidelines in the literature (*7*).

### Recordings

We assessed the quality of the recordings of the NeuroRoots after implantation targeted to the CA1 of the hippocampus. The raw signal exhibited a high signal-to-noise ratio (SNR) of 4.1 that allowed for clear identification of action potentials (APs) across the different channels (Fig. 3.a.i). Although large local field potentials (LFPs) could be recorded across multiple adjacent electrodes, we could selectively record localized APs on different channels without cross-talks (Fig. 3.a.ii).

**Figure 3:**
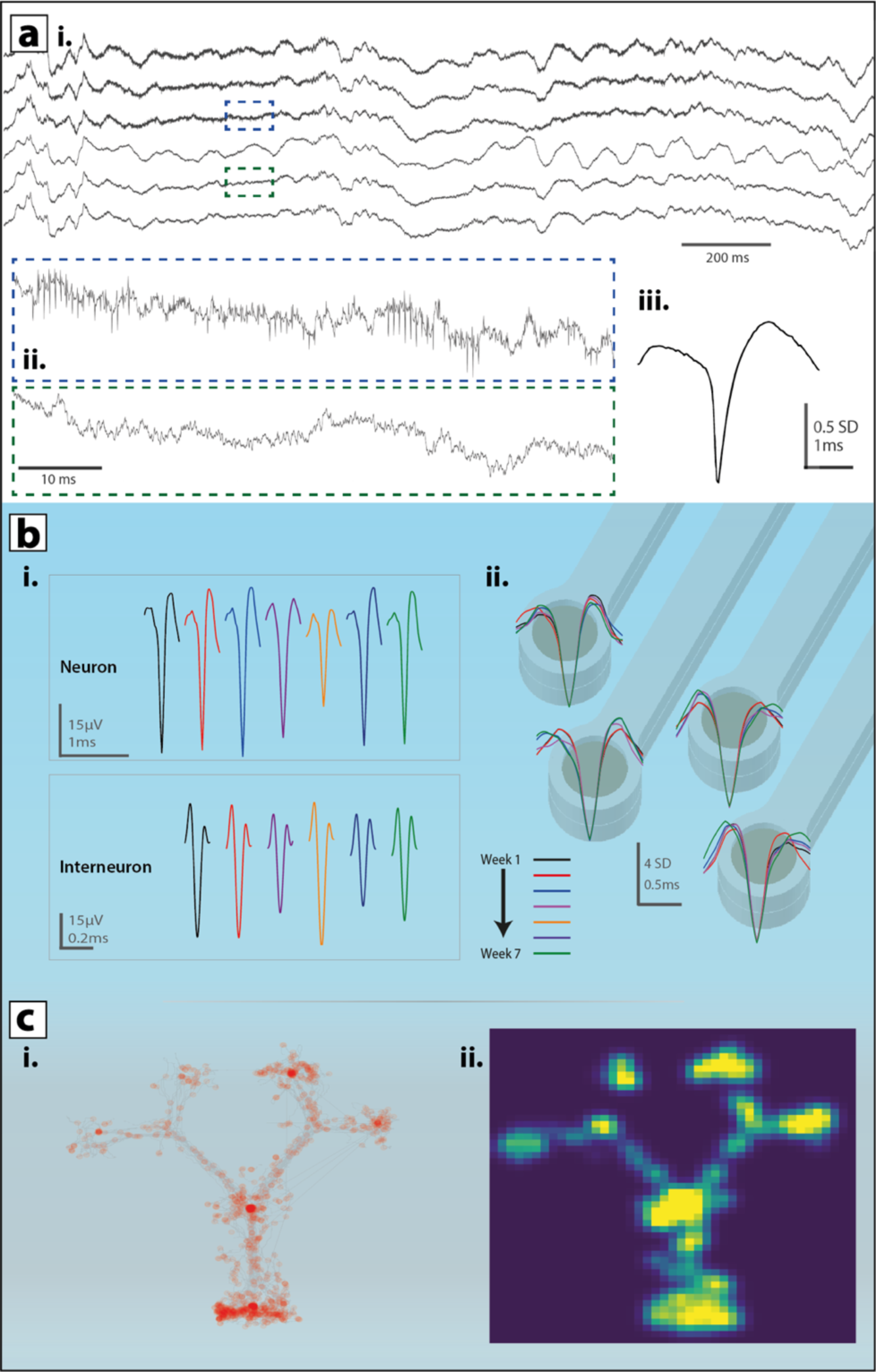
**a) Representative recordings of raw traces.** i) Two seconds recording of 6 consecutive channels. ii) Zoomed-in view of the blue box presented in i) shows characteristic activity of an interneuron burst while the green zoomed-in green box show the activity of a neighbor electrode. ii) action potential of a hippocampal neuron recorded during the acute experiment. **b) Cluster stability assessment during the chronic experiment.** i) Representative APs of the same neuron and interneuron over the 7 weeks of the experiment. ii) Overlay of averaged APs corresponding to a cluster tracked over 7 weeks. Each color corresponds to the averaged data over 10 minutes recording. Each cluster corresponds to a different electrode. Each color is representative of recordings from a different week as describes in the colored legend. **c) Behavioral analysis**. i) animal trajectories in grey on a double Y-Maze (1.4m x 1.2m). Spikes from an example cell overlaid in red. ii) estimated firing fields for the same cell, warmer color indicates increased firing activity.

The averaging of the signal into bins of 1 ms centered around sortable spikes revealed a variety of distinct extracellular spike waveforms, spread along 32 channels of the NeuroRoots probe (Fig. 3 a-b). Analysis of the AP waveforms could distinguish between two distinct types of activity, interneurons characterized by a fast ∼0.3 ms spike of 60 µV peak-to-valley amplitude and neurons of ∼1 ms spike and 50 µV amplitude (Fig. 3.b.i).

We then evaluated the stability of the electrical coupling of the NeuroRoots with the brain by recording the neuronal activity across an implantation period of seven weeks without any further adjustment of the electrodes. The analysis of all 32 channels chronically implanted into the HIP while the rat was freely moving highlights several points.

First, APs could be observed a few minutes only after the surgery, suggesting the damage done during the electrode implantation were low enough to preserve spontaneous activity in the neural tissue around the probe (Fig. 3.a.iii), and thus are recording from pre-existing neural circuits. This is supported by the short recovery period needed to start freely-moving chronic recordings after the surgery as sortable APs could be observed on all channels during our first experiment 4 days post-surgery.

Second, the similarity of extracellular action potentials over the time of the experiment suggest that the NeuroRoots have the ability to form a stable interface with the surrounding neurons (Fig. 3.b.ii). Moreover, the interneuron activity was also included in the analysis as a high-probability marker for tracking the same neuron as previously reported. While the shape of the AP was highly consistent, the absolute magnitude varied from week to week (Fig. 3.b.i). This variation was non-monotonic, sometimes increasing or decreasing over time, suggesting it could result from natural remodeling near the neuron or evolution in local cell architecture (*36*).

In order to assess whether these were likely to be the same neurons over time, we performed cell sorting using unsupervised clustering and Principal Component Analysis (PCA) (Fig. S3). The result of that analysis over the seven weeks showed minimal shifts of the cluster center on the same electrode, equivalent to 0.61 σ between the first and last dates. Together with the consistent signal shape between the different dates, this suggests that the waveforms were generated by the same neuron. Analysis of the signal to noise ratio (SNR) for all 32 channels between the first last recording dates showed a remarkable stability, with less of 3.1% variation around the average. This indicates that the firing neurons remained in close proximity to the electrode for the entire duration, and that the electrode did not undergo any detectable degradation.

Since the device could be used for freely moving animals, we monitored the position of the rat moving within a double Y-Maze and compiled a comparative map of positions versus firing rate (Fig. 3.c). The animal trajectories are represented in grey and the spikes from an example cell overlaid in red. Spatial firing fields were estimated for the same cells, which allows to determine if the cell exhibited any spatial selectivity. The maximum firing rate recorded was 1.02 spikes/s, which although too low to indicate the monitoring of a cell selective for position, demonstrates the compatibility of the NeuroRoots platform with measurements of interest in behavioral experiments.

## Discussion

NeuroRoots combines several key innovations toward the development of integrative BMI implants with simple delivery strategy, stable recordings and scalable number of electrodes.

### High density electrode distribution can be tailored to match the application

Unlike conventional shank electrodes, the NeuroRoots recording pad distribution does not have to be uniform and could for instance have several regions of ultra-dense sampling up to ∼5x the current state of the art (*17*), or a sparse sampling over large distances. This could be advantageous for studying local neural architecture (*37*, *38*) or dynamics and plasticity in behaving animals under different brain or behavioral states (*39*). The implantation of a high electrode density with minimal disruption of existing neural circuits also opens the prospect of electrical interfacing with neurons located into dense cellular regions of the brain as well as simultaneous multiple-site implantations.

### Scalable design allows for high channel count with minimal footprint

We demonstrated a bio-integrative approach to implant electrodes in brain tissue using a few dozen channels, which was largely limited by the currently available connectors. As new connectorization technologies emerge, the unique form factor of NeuroRoots will allow for scaling up the number of channels to a few hundred without dramatically increasing the implant footprint. For example, a 10-fold increase in channel count (320 electrodes) increases the diameter of the implant by a factor of two (60µm in diameter), which is still quite small compared to current Michigan or Utah style devices. We anticipate that this dramatic increase in channel count will open new possibilities in research and clinical applications for BMIs.

### Minimal footprint, ultra-flexibility enables recording stability

The ultra-flexible leads and open-geometry of the NeuroRoots offer well matched mechanical compliance to brain tissue and allow for tissue ingrowth around the implant, potentially accounting for the improved recording stability. The small electrical lead dimensions (∼5 µm wide) also allow rapid diffusion of oxygen and signaling molecules around the device, which could also contribute to minimizing immunogenicity. For example, a 5 µm wide electrode positioned between two neurons spaced 100 µm apart is predicted to increase diffusion times just 0.1%, indicating the implant itself should not interfere with natural communication. This ability to allow for inter diffusion of signaling molecules in and around the implant is important for long term stability as disrupting cellular communication can trigger the foreign body response (*40*) (refs).

### Compatibility with existing equipment facilitates widespread adaptation

Many innovative BMI devices have struggled to gain significant traction as either research tools or clinical devices due to complicated implantation methods and/or incompatibility with established infrastructure including surgical tools and recording equipment. In contrast, the NeuroRoots was specifically designed to be easy to implant and to be readily integrated with existing equipment. This compatibility has already been proven in two distinct research facilities (Stanford and Aix-Marseille) that use standard connectors and recording equipment (Neuralynx and Intan). The NeuroRoots’ inherent mechanical flexibility also reduces the potential for the type of mechanical failures that plague silicon based neural probes. Looking ahead, we foresee that these advantages will significantly reduce the barriers to widespread adaptation of NeuroRoots by neuroscientists and clinical partners.

### Conclusion

In this work, we introduce bio-mimetic NeuroRoots electrodes with similar size, flexibility and distribution as axon bundles in the brain. We demonstrate the implantation of a dense distribution of 32 of these electrodes into the brain of freely moving rats with minimal acute damage and long-term stable integration within the tissue. Stable neural recordings were made over seven weeks during complex behavioral tasks with freely-moving rats. The combination of scalability, low damage, stable single unit recording, and ready integration with existing surgical and recording equipment make NeuroRoots a promising candidate for basic neuroscience experiments and clinical applications.

## Materials and Methods

### Probe fabrication and preparation

The fabrication and patterning of Parylene C and PEDOT:PSS based electrodes were discussed in previous publications(*2*, *41*), resulting in devices capable of conformability around a 35 µm diameter microwire (Fig. 1.f). Here we used an adapted fabrication process consisting of deposition and patterning of parylene C, and Pt as follows: Parylene C was deposited using an SCS Labcoater 2 to a thickness of 1.5 µm (to ensure pinhole-free films). 3-(trimethoxysilyl)propyl methacrylate (A-174 Silane) and a dilute solution of industrial cleaner (Micro-90) were used as an adhesion promoter and anti-adhesion, respectively. The film was patterned with a 150 µm thick layer of Germanium and dry etched by a plasma reactive-ion etching process (500 W, 50 sccm O2, for 5 min) using P5000 followed by an immersion into deionized water in order to dissolve the metal mask. A dual layer resist liftoff process was used to pattern metal pads and interconnects. A first resist, Shipley LOL2000, was spin-coated on the Parylene C film at 5,000 r.p.m., baked at 200 °C for 15 minutes. A positive photoresist, Shipley 955 i-line then spin-coated at 3000rpm, baked at 110 °C for 90 seconds and then exposed using an ASML stepper (ASML PAS 550 0/60 i-line Stepper), and then developed using MF26A developer. Metallic layers (10 nm Ti, 150 nm Pt) were deposited using an e-beam metal evaporator (Innotec ES26C) at 2.10-6 bars. Lift-off was performed using 1165 stripper (2 hours). For the devices with PEDOT:PSS coatings, the basic fabrication process followed previously reported procedures including a Parylene C peel-off step to pattern the PEDOT:PSS (*41*).

The electrodes were characterized in vitro using Phosphate Buffer Solution (PBS) solution. An Ag/AgCl wire was immersed in the electrolyte and used as the reference electrode during impedance measurements.

### Shuttle microwire preparation and assembly with guiding system

The microwire was prepared using a technique previously reported (*42*). A 2M KOH was prepared by dissolution of KOH dices (Fischer) into deionized water. A 35µm diameter tungsten wire (Goodfellow USA) was slide into a 100µm inner-diameter polyimide tubing (Neuralynx) leaving several centimeters protruding on each side. The microwire was then electrosharpened using a 2V DC bias against an Ag/AgCl reference electrode. The protruding length of the microwire was then adjusted to 5 mm to up to 1 cm and the other extremity was sealed to the polyimide tubing to prevent sliding of the microwire. The NeuroRoots were then connected to the ZIF connector and assembled using the technique described previously in this paper.

### Animals surgery for chronic recordings

Two adult Long Evans male rats aged 3-4 months and weighing 400-500g were used in this study (Charles River Laboratories). Animals were singly housed under a regular 12hr light/dark cycle, with experiments carried out during the light cycle. Standard surgery procedures were followed using a stereotaxic platform. To target the CA1 region of the hippocampus we performed a 2.5 × 2.5mm craniotomy (−3.6 mm AP, −2.2 mm ML from Bregma). All procedures and animal care were approved by the Institutional Animal Care and Use Committee at Stanford University School of Medicine.

In typical procedures, the dura mater was removed, and the implant inserted into the brain using a stereotaxic frame. The device mounted into the 3D printed hat was vertically mounted on a micromanipulator ((Model 963, Kopf Instruments) and positioned above the craniotomy hole. As the device traveled downward, the protruding tip microwire/NeuroRoots penetrates the neural tissue. Once the neural probe reached the desired depth and the coated dissolution time achieved, the shuttle microwire was retracted, and the NeuroRoots was released and left embedded in the brain tissue. Because of the minimal footprint of the NeuroRoots, the craniotomy was kept minimal with a diameter of ∼2-3mm in order to keep the surgery least invasive as possible. The exposed surrounding tissue was covered with Kwik-Sil (World Precision Instruments), and the hat was secured to the rodent’s skull using standard procedures with initial layers of Metabond and dental cement.

### µCT imaging

Three dimensional computerized X-ray tomography images were performed to image NeuroRoots devices with PEDOT:PSS coated electrodes implanted 3 mm deep into a rat brain. Implantation of the NeuroRoots device was performed immediately following the extraction of the brain of a rat using a 100 µm diameter microwire as the shuttle. The sample was then immersed in fixative solution (2% formaldehyde) for 6 days before imaging. Images were taken using a Zeiss Versa 510 (80 kV excitation voltage, 7 W power). Image processing was done with the associated Zeiss software and care was taken to ensure feature dimensions in the uCT images were consistent with measurements from optical microscopy.

### Histological sample preparation

Animals were transcardially perfused first with saline, then with 150 ml of fixative solution containing 4% PFA in 0.1 M phosphate buffer (PB). Tissue blocks were cut horizontally on a Vibratome (Leica VT1200S, Leica Microsystems, France) into 40 µm sections After extensive washes in PB, GFAP staining was used (GFAP Monoclonal Antibody (GA5), Alexa Fluor 488, Thermofisher, France). Sections were mounted on SuperFrost slides and covered with a mounting medium containing 2-(4-amidinophenyl)-1H-indole-6-carboxamidine (DAPI) (Flouromount Mounting Medium with DAPI, Abcam, UK).

### Data acquisition and processing

Data were collected using a Digital Lynx SX acquisition system (Neuralynx Inc). Local field potential signals were collected for the 32 channels and locally amplified with an active headstage device (HS-72-QC, Neuralynx Inc.) that magnetically attaches to the NeuroRoots implant. Signals were sampled at 32kHz. The headstage further provided positional information through mounted LEDs that were tracked via an overhead camera.

Rat LFP and unit activity were recorded either in open field of in a double Y-Maze (1.4m x 1.2m) during normal behavior or trained tasks. The data were analyzed using MATLAB (MathWorks).

Spike detection was achieved through the following processing steps: filtering of the data with a bandpass filter set to 600Hz –7000Hz, Principal Component Analysis (PCA) and k-means clustering. From the threshold analysis of each individual channel, 1 ms events were extracted, centered around each peak. The PCA features were then computed, and the data was projected onto the 10 largest components to generate the feature vectors. Subsequently, k-means clustering, an unsupervised learning method, was applied, with k ranging from 2 to 4. Through manual curation, each cluster was consolidated over time. The 2 largest PCA components of the first recorded date of the cluster were used as the basis vectors. For each date, the cluster points were projected onto the basis vectors, and a multivariate Gaussian distribution was subsequently fitted. We used the mahalonobis distance of the mean of the distribution of the week 7 to the distribution of week 1 to calculate the variance of the PCA centers.

The signal-to-noise ratio (SNR) for each channel was computed by dividing the average spike amplitude by its corresponding noise level. The noise level is estimated as the median(|V|)/0.6745.

## Acknowledgments

**General:** The authors thank A. Obaid (Stanford University) and D. Khodagholy and J.N. Gelinas (Columbia University) for fruitful discussions. We also thank Makoto Nakamura from ASML and the Stanford Nanofabrication Facility (SNF) for help with the device fabrication, Sawson Taheri from the Stanford Prototyping Facility (SPF) for the help with the PCB design, Quentin Payan from WurthElektronik for the ZIF connectors and the Biosphera team for the original rat 3D model used in Figure 2.b. We thank Damiano Barone and Anthony Dennis respectively for assistance with µCT sample preparation and µCT imaging.

**Funding:** M.F. was supported by US National Institutes of Health Grants (1R21EY026365-01) and the Stanford NeuroTechnology Initiative (NTI). C.P. acknowledges funding from a Whitaker International Scholar Grant administered by the U.S. Institute for International Education as well as funding from the University of Cambridge Borysiewicz Biomedical Fellowship program. A.W. acknowledges funding from the European Research Council (ERC) under the European Union’s Horizon 2020 research and innovation programme (grant agreement No 716867).

**Author contributions:** M.F. and C.P. were primary experimental contributors. M.F, C.P, N.M conceived the basic device concept and designed the experiments. M.F. designed and fabricated the devices and platform for the chronic recording experiments. A.G and M.F recorded the chronic signals and M.F, A.G and E.Z analyzed the electrophysiological data. C.P., M.F., J.P., G.D., M.D., fabricated and characterized the devices for acute and histology experiments. A.S. performed histology preparation and imaging. M.F. and N.M wrote the manuscript with input from C.P.. All authors reviewed the manuscript and provided comments.

**Competing interests:** The authors declare no competing interests.

**Data and materials availability:** All data needed to evaluate the conclusions in the paper are present in the paper and/or the Supplementary Materials. Additional data related to this paper may be requested from the authors.

## Supplementary Materials

**Fig. S1.**
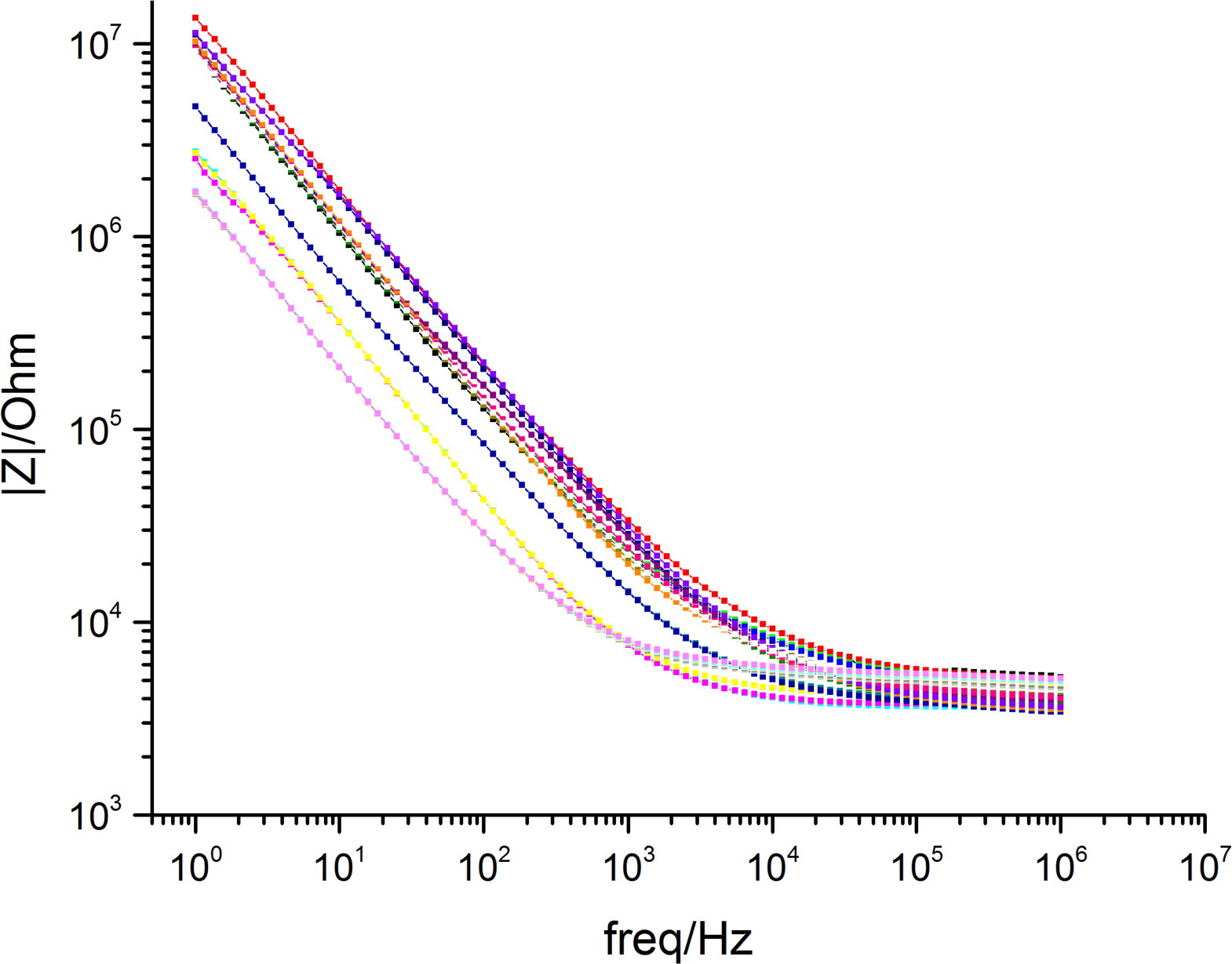
Impedance of NeuroRoots electrodes

**Fig. S2.**
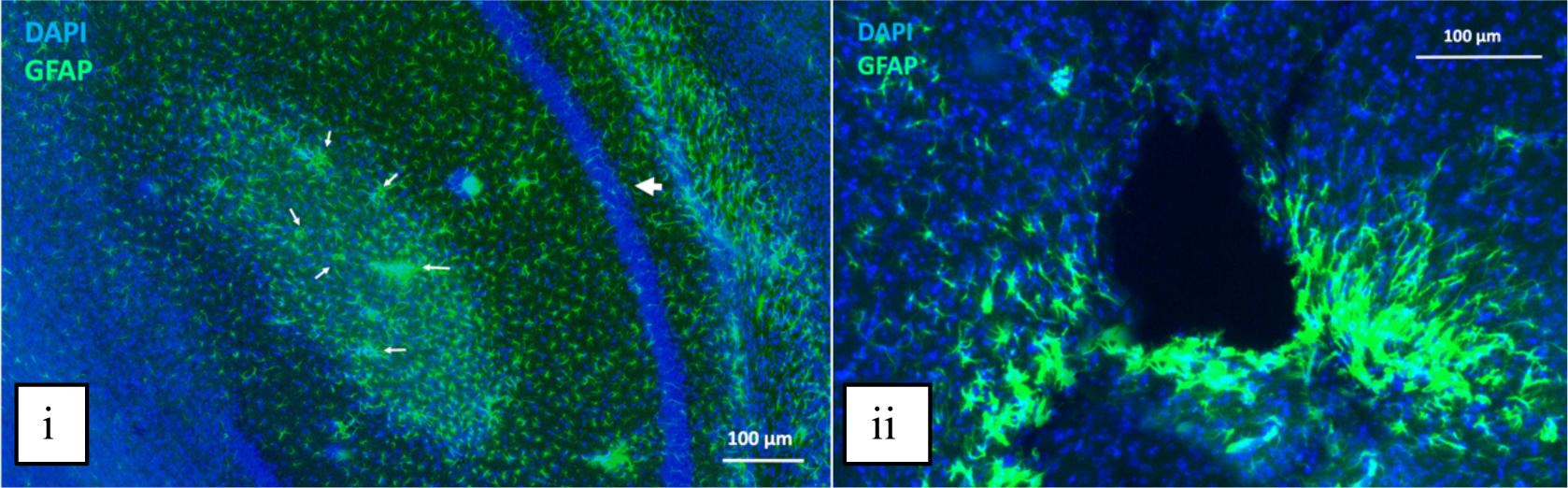
Histological evaluation of NeuroRoot along the insertion trajectory. Histological verification of a 90-days-post-implantated, large PEDOT:PSS coated NeuroRoots implanted with a 100 µm diameter microwire. High magnification images show horizontal cross sections of the tissue response around the (i) tip of the implant and (ii) along the trajectory of the implant. GFAP staining (green) show reactive astrocytes 90 days post-implantation whereas DAPI (blue) labels cell nuclei in the neural tissue. Scale bar: 100 µm

**Fig. S3.**
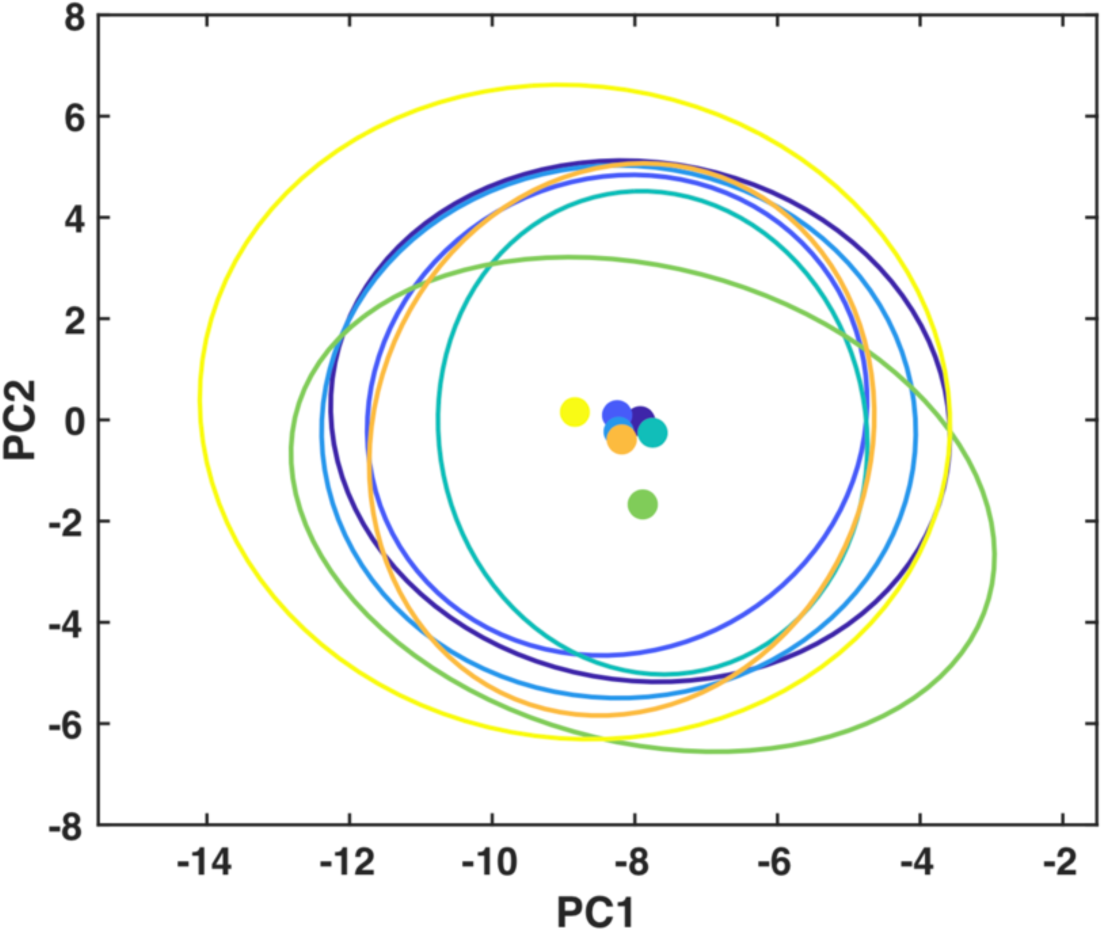
Principal Component Analysis of one electrode cluster

